# HCN2 in cholinergic interneurons of the nucleus accumbens mediates reward response

**DOI:** 10.1101/2021.09.21.460269

**Authors:** J Lee, M Weinberger, Y Kawahara, J Cheng, G Umschweif, L Medrihan, M Flajolet, A Nishi, Y Sagi

## Abstract

Cholinergic interneurons (ChIs) of the nucleus accumbens (NAc) are important for mediating the behavioral response to rewarding stimuli. A major role for these cells is to regulate dopamine (DA) transmission by activating cholinergic receptors at local DAergic nerve terminals. However, the mechanisms that enable cholinergic neurons to enhance DA release in response to reward remain unknown. Here we report that the hyperpolarization-activated cyclic nucleotide-gated channel 2 (HCN2) in NAc ChIs mediates an enhancement in DA signaling in response to rewarding stimuli. The HCN current in NAc ChIs and its modulation by DA, as well as the increase in cholinergic efflux by local cocaine infusion were impaired in mice with deletion of HCN2 in cholinergic cells. Enhancement in the DA efflux and signaling in the NAc in response to rewarding stimuli, as well as cocaine conditioning were also dependent on HCN2 in ChIs. These results provide a mechanistic link between the activity of NAc ChIs and reward encoding.

## Introduction

The mesolimbic projection pathway which controls dopamine (DA) transmission in the nucleus accumbens (NAc), plays a central role in mediating reward encoding in response to rewarding stimuli. In turn, a dysfunction in this pathway is implicated in mental disorders, including drug abuse and depression [1]. Among the different cell populations in the NAc, cholinergic interneurons (ChIs) are unique because they directly regulate synaptic DA release [2].

Acetylcholine released from NAc ChIs activates presynaptic nicotinic receptors at DAergic nerve terminals, leading to enhanced DAergic signaling [3]. The idea that the activity of ChIs enhances reward encoding is supported by functional studies. Inhibition of NAc ChIs impaired motivation for cocaine-induced conditioned place preference (CPP) [4], and inhibition of NAc ChIs activity or transmission induced anhedonia-like behavior [5, 6]. Additionally, the levels of acetylcholine and DA in the NAc were up-regulated by rewarding stimuli [7, 8], whereas inhibition or activation of NAc ChIs respectively attenuated or enhanced the efflux of DA in response to local cocaine infusion [8].

Autonomous tonic cell firing activity of ChIs is regulated by an ensemble of ion channels, including the hyperpolarization-activated cyclic nucleotide-gated (HCN) channel family [9-11]. A molecular link was recently found between HCN channels in NAc ChIs and reward encoding. Anhedonia-like behavior in mice was mediated by attenuation of the firing rate of NAc ChIs [5]. Notably, the attenuation in the tonic activity in NAc ChIs in the depressed-like animals was mediated by down-regulation of HCN2 in NAc ChIs [5]. To better understand the link between reward and the function of NAc ChIs, we evaluated the role of HCN2 in regulating changes in the function of NAc ChIs by DA and reward.

## Materials and Method

### Animals

All procedures were approved by the Animal Care and Use Committee of the Rockefeller University. All animals were maintained on a 12-hour light/ dark interval with food and water *ad-libitum*. Transgenic mice were backcrossed with C57/ Bl6N background mice for at least 6 generations. Male mice between 8 to 12 weeks of age were used for behavioral tests. Male and female mice were used for molecular and physiological studies, except for DA microdialysis, with no differences observed between genders. For social defeat, CD-1 male retired breeders were purchased from Charles River Laboratories. Mice with a floxed HCN2 allele were generated by introducing two loxP sites around exon 2 of *hcn2* (NM_008226.2). To validate its deletion in tissues HCN2 ^*fl/fl*^ mice were crossed with Nestin ^*Cr+/-*^ (B6.Cg(SJL)-TN (NesCre)1Kln/J) mice. In all studies that compared WT and cKO, WT mice (ChAT ^*Cre* -/-^) and cKO (ChAT ^*Cre* +/-^) were littermates crossed with HCN ^*fl/fl*^. For studies that included immunohistochemistry or electrophysiology, animals were crossed with ChAT ^*EGFP*^ (GH293) accordingly, resulting in WT, ChAT ^*Cre* -/-^ :: ChAT ^*EGFP*^:: HCN ^*fl/fl*^, and cKO, ChAT ^*Cre* +/-^ :: ChAT ^*EGFP*^:: HCN ^*fl/fl*^.

### HCN2 gene expression and protein levels analysis

mRNA isolation, purification, cDNA amplification and qPCR analysis of mRNA levels were as previously described [12, 13]. For protein purification, NAc was punched from freshly dissected mouse and was lysed in 1% SDS. Protein concentration was determined using the BCA protein assay (Thermo Fisher Scientific, Waltham, MA). 20 μg of protein were loaded onto 4-12% Bis-TRIS gels and transferred to a PVDF membrane for western Blotting analysis. Proteins were detected using antibodies for HCN2 (14957) and β-Actin (4970, Cell Signaling Inc., Danvers, MA).

### Immunohistochemistry

GFP and HCN2 were detected using antibodies (1218 from Abcam and 14957 from Cell Signaling, respectively). For detection, secondary Alexa goat anti-mouse or goat anti-rabbit were used (Thermo Fisher Scientific, Waltham, MA). Auto fluorescence was used to detect mCherry. Cell nuclei were detected using DRAQ5 (Thermo Fisher Scientific). To quantify HCN2 expressing cells, 3-4 coronal sections of 45 μm thickness were stained per mouse. The pixel mean value of HCN2 inside a GFP positive ChI was divided by its level outside the cell, using a custom written Matlab code ^5^. Cells with HCN2 mean pixel values above 140% of the background were considered immunopositive.

### Electrophysiological Recording

#### NAc slice preparation and electrophysiology

3 to 4 weeks old mice were euthanized with CO_2_, their brains were quickly removed and placed in an ice-cold N-Methyl-D-glucamine (NMDG)-containing cutting solution (in mM: 93 NMDG, 2.5 KCl, 1.2 NaH_2_PO_4_, 30 NaHCO_3_, 25 glucose, 20 HEPES, 5 sodium ascorbate, 3 sodium pyruvate, 2 thiourea, 0.5 CaCl_2_, 10 MgSO_4_, pH 7.4, 295-305 mOsm). Coronal brain sections (300 μm thickness) containing the NAc were sliced using a VT1000 S Vibratome (Leica Microsystems Inc., Buffalo Grove, IL, USA). After sectioning, slices were allowed to recover in the cutting solution saturated with 95% O_2_ and 5% CO_2_ for 15 min at 37 °C. The slices were then transferred and incubated in the recording solution at room temperature for at least 1 hour before recording.

NAc-containing slices were placed in a perfusion chamber attached to the fixed stage of an upright BX51WI microscope (Olympus, Japan) and submerged in continuously flowing oxygenated recording solution containing (in mM: 125 NaCl, 25 NaHCO_3_, 25 glucose, 2.5 KCl, 1.25 NaH_2_PO_4_, 2 CaCl_2_, and 1 MgCl_2_, pH 7.4, 295-305 mOsm). Neurons were visualized with a 40x water immersion lens and illuminated with near infrared (IR) light. Electrophysiological recordings were made with a Multiclamp 700B/ Digidata1550 system (Molecular Devices, Sunnyvale, CA, USA). Patch electrodes were made by pulling TW150-4 glass capillaries (World Precision Instruments, Sarasota, FL, USA) on a PP-830 Single Stage Glass Microelectrode Puller (Narishige, East Meadow, NY, USA). The pipette resistance was typically 3-5 MΩ after filling with the internal solution (in mM: 126 K-gluconate, 10 KCl, 2 MgSO_4_, 0.1 BAPTA, 10 HEPES, 4 ATP, 0.3 GTP and 10 phosphocreatine, pH 7.3, 290 mOsmol). All electrophysiological recordings were carried out blind to the experimental conditions including genotype and drug treatment.

Whole-cell voltage-clamp was made from the soma of ChIs to measure HCN currents, voltage-gated potassium currents and voltage-gated sodium currents. HCN currents were recorded with a series of three second pulses with 10 mV command voltage steps from -150 mV to -70 mV from a holding potential at -70 mV. Tetradotoxin (TTX, 1 μM) and tetraethyl ammonium chloride (TEA, 5 mM) were added in the aCSF (pH 7.4, 295-305 mOsm) to block sodium currents and potassium currents, respectively. HCN current amplitudes were calculated as the difference between the instantaneous current at the beginning of the voltage step and the steady-state current at the end of the voltage step [14].

All data were acquired at a sampling frequency of 50 kHz, filtered at 1 kHz and analyzed using pClamp10 software (Molecular Devices, San Jose, CA).

### Microdialysis

All stereotaxic surgeries were performed on an Angle Two Small Animal Stereotaxic Instrument (Leica Biosystems Inc., Buffalo Grove, IL, USA). Mice were anesthetized by intraperitoneal ketamine/ xylazine (100/ 8 mg/ kg) and a unilateral guide cannula (CXSG-4, Amuza Inc., CA) was placed in the NAc. The coordinates for the cannula tip were +1.30, +1.10, -3.15 (L, A, V) mm from Bregma. The guide cannula was mounted to the skull using Metabond cement (Parkell, Edgewood, NY), and a dummy probe was placed. After recovery of 7± 1.3 days, the animals were briefly anesthetized using 1.5% isoflurane. 2 mm long concentric probe with outer diameter of 0.22 mm, was slowly lowered to extend directly from the cannula tip (CX-I-4-2, Amuza, CA). The animals were then returned to its home cage with free access to food and water. Dialysis started using solutions that were warmed to 35-37ºc immediately before use. After the placement of the probe, aCSF (Harvard Apparatus, MA) was dialyzed at 0.05 μl/ min for two hours, and then changed to 1 μl/ min for 30 min after which dialysate fractions were automatically collected (BASi, ME). At the end of the dialysis session animals were immediately euthanized and the position of the probe in the NAc was visually confirmed. ACh content was detected using HPLC-ECD system with AC-GEL separation column (2.0 ID x 150 mm) with a platinum working electrode (all from Amuza, San Diego, CA). The applied potential was 450 mV and the background current was 3.7-9 nA for all samples. Mobile phase consisted of 50 mM KHCO3, 135 μM Na_2_-EDTA, and 1.6 mM sodium 1-decanesulfonate obtained retention time of 13.71 ± 0.021 min. ACh content in each dialysate sample was determined using subsequent standards with known amounts of ACh. The threshold for detection was 2.84 fmol/ min ACh. For each mouse, 24 samples of 3-minute long each were collected. 4 without and 8 with neostigmine (500 nM). The solution was then replaced and cocaine (1μM) was added and 12 more samples were collected. Data from mice with ACh levels above threshold in all of their 9^th^ – 12^th^ dialysates were included for analysis. For these mice, respective averages of ACh content from fractions 1-4^th^, 9^th^ -12^th^ and 21^st^ -24^th^ were used for statistical analysis.

Detection of dopamine by microdialysis was done as previously described [8]. Briefly, microdialysis probes (I-shaped cannulas) were implanted unilaterally in the NAc (exposed length 1.5 mm) of mice with real-time quantification of dopamine. Ringer’s solution without neostigmine was used for dialysis at a flow rate of 2.0 µl/ min. The 20-min sample fractions collected through the dialysis probes were directly injected to high-performance liquid chromatography using a reverse-phase column (150×4.6 mm, Supelcosil LC18; Merck, Darmstadt, Germany) with electrochemical detection. An LC-20AD pump (Shimadzu Corporation, Kyoto, Japan) was used in conjunction with an electrochemical detector (potential of the first cell, +180 mV; potential of the second cell, -180 mV) (ESA, Chelmsford, MA). The mobile phase was a mixture of 4.1 g/ L sodium acetate adjusted to pH 4.1, 100 mg/ L Na2EDTA, 120 mg/ L octanesulfonic acid and 10% methanol. The flow rate was 0.4 ml/ min. The detection limit of the assay was approximately 0.9 fmol per sample (on-column).

### Photometric detection of dopamine signaling

Mice were injected with AAV5 expressing dLight 1.3b to the medial NAc. Coordinates were: +1.45, +1.20, -4.61 mm (L, A, V) relative to Bregma, respectively. After two weeks, 0.4 mm O.D cannulated silica fiber (Doric Inc., Quebec, Canada) was implanted and its ferule was secured to the skull using Metabond (Parkell Inc., Edgewood, NY). Each mouse was recorded once, 7± 0.4 days after cannulation. The animal was anesthetized using 1.5% isoflurane, and placed on heating pad set to 37°C. Head stage with a light source of excitation wavelength LED peak 470 nm, filter band 445∼490 nm and a detector unit with emission wavelength filter band 500∼550 nm and sampling rate of 100 Hz and a wireless transmitter was secured to the ferule, and photometric traces with temporal previsions of 10 msec and time stamps of drug injections were recorded and filtered for 5 Hz online (Amuza, CA). The Excitation power was ∼300 µW. To minimize variability in photobleaching between animals, the power was adjusted at the beginning of each session, keeping the initial baseline signal amplitude at 100 ± 0.8% for all animals. Cocaine hydrochloride (1.5 mg/ kg), donepezil hydrochloride (0.2 or 2 mg/ kg) and nicotine hydrogen tartrate (0.07 mg/ kg) were freshly dissolved in saline right before they were injected intraperitoneally. After the recording, the animals were euthanized by transcardial perfusion and the localization of the fiber and the presence of GFP transgene in the NAc were validated by immunohistochemistry. Curves of fluorescence decay were calculated offline using quadratic regression and curves with R^2^ > 0.95 were used as a signal control. ΔF/ F values were determined for traces of 15 s (5 s before the injection and 10 s after) using pMAT1.2 [15]. ΔF/ F peak amplitude and AUC values were determined by GraphPad Prism 9.2. Amplitude and AUC values within 2 – 8 s post-injection for saline, cocaine and donepezil, or 4 – 10 s post-injection for nicotine, above a threshold of 25% of the maximum were included in the statistical analysis.

### Behavioral Tests

Behavioral tests were conducted and analyzed by a researcher blind to the genotype.

### Open Field

Mice freely explored an arena (50 cm x 50 cm x 22.5 cm) during 60 min, immediately after cocaine (15 mg/ kg) or saline injection. The total distance traveled was automatically recorded using the automated Superflex software (Accuscan Instruments, Columbus, OH, USA). Distance traveled during each 10 min segment was plotted and used for statistical analysis.

### Social Approach Test

The subject mouse was allowed to freely explore a 3-chamber-arena (58.4 cm x 42.6 cm x 23.1 cm) containing two empty wire mesh cups for 10 minutes. Then, an unfamiliar mouse (male of the same age and fur color) was placed inside a cup in one chamber, and a Lego object was placed inside another identical cup in the opposite compartment. The subject mouse was placed in the middle compartment and allowed to freely explore the cups for 10 min. Video tracking software (Ethovision XT7, Noldus, VA) was used to record the session. Social approach was defined by contact between the snout of the subject mouse and the wire mesh cup. The total duration as well as the number of interactions (visits) with the unfamiliar mouse and the object were scored offline.

### Sub-threshold defeat and social interaction

The sub-threshold social defeat stress paradigm was carried out as described previously [16], with mild modifications. Mice underwent 3 defeat sessions in which they were placed into the home cage of a CD-1 aggressor for 3 min of physical contact, and returned to their home cage to rest for 15 min. After the last defeat session, the experimental mouse was returned to a group housing in its home cage. Social interaction test was performed next day. The test was composed of two phases, 2.5 min each, in which the experimental mouse was allowed to explore an open field arena (42 cm x 42 cm x42 cm) with a wire mesh enclosure (10 cm wide x 6.5 cm deep x 42 cm high) located in a designated place inside the arena. In the first phase, the enclosure was empty. In the second phase, a novel CD-1 aggressor mouse was placed inside the enclosure. The amount of time spent by the experimental mouse in the interaction zone (IZ) surrounding the wire mesh was recorded and analyzed by the video-tracking apparatus and software Ethnovision 7.0 (Noldus, Wageningen, the Netherlands). Interaction ratio was calculated by dividing the amount of time spent by the experimental mouse in the IZ in the second phase over the time in the IZ in the first phase.

### Place conditioning and chemogenetics

Mice were placed in a 3-chambered apparatus (cocaine/ saline chambers 16.8 × 12.7 × 12.7 cm, connected by a small chamber, Med Associates, Columbus, OH), in which each chamber was distinguished by both wall color and floor texture. Training and testing were performed at the same time of the day. During testing, animals were placed in the central compartment and exposed to the entire apparatus for 15 min. Time spent and number of beam breaks in each chamber were automatically recorded using CPP apparatus 3013 (Med Associates, Fairfax, VT), and was used for statistical analysis. On the first day, mice were given a pre-test and the chamber with lesser time spent was paired with cocaine. Conditioning training was on days 2-9 and extinction training on days 11-18. Home cage abstinence was on days 22-50. Acquisition test was on day 10, extinction tests on days 19, 21, 51 and 53, and reinstatement tests on days 20 and 52. During conditioning and extinction training, mice were confined inside the paired-compartment for 30 min. Saline (100 µl) or cocaine (15 mg/ kg) were injected intraperitoneally immediately before conditioning training, and cocaine immediately before reinstatement tests. For studies with viral injections, behavioral handling started 3 weeks after the surgery. 500 nl of AAV expressing Gi-DREADD, Gs-DREADD or mCherry or AAV2-FLEX-HCN2 [5] or its control AAV-FLEX-GFP were injected into the NAc of mice using a 10 μl Hamilton syringe at a speed of 0.1 μl/ min. DREADD viruses were injected to HCN2 ^*WT/ WT*^:: ChAT ^*Cre+/-*^ :: ChAT ^*EGFP*^ mice. HCN2 or GFP O/E AAV were injected to HCN2 ^*WT/ WT*^:: ChAT ^*Cre+/-*^ or cKO mice. Coordinates were: ±1.45, +1.20, -4.61 mm (L, A, V) relative to Bregma, respectively. The needle was left for additional 10 min and then was slowly withdrawn. The stereotaxic injections were confirmed by immunohistochemistry to adequately cover the NAc without significant spread to other regions. For chemogenetic manipulation, Clozapine N-oxide (CNO, 4 mg/ kg, C0832, Millipore Sigma, St. Louis, MO) was dissolved in saline and was injected 15 min before cocaine.

### Statistical analysis

Unless mentioned otherwise, bar graphs and dot plots are presented as means ± s.e.m. Sample size was chosen based on previous reports to ensure adequate statistical power. Statistical analysis was performed using Prism 9.2 (GraphPad Software, La Jolla, CA, USA), or SAS MIMED (version 9.4, SAS Institute, Cary, NC). In all experiments, P< 0.05 was considered significant.

## Results

### Cholinergic response to dopamine signaling in the NAc is attenuated by deletion of HCN2 in cholinergic interneurons

Since HCN2 might regulate ChIs activity, to identify the functional role of HCN2 in these cells, we generated a transgenic mouse model bearing floxed HCN2 allele. Following crossing HCN ^*fl/fl*^ with either Nestin ^*Cre*^ or ChAT ^*Cre*^ mice, we noticed the loss of HCN2 mRNA and protein expression in brain tissue from the Nestin ^*Cre*^ :: HCN ^*fl/fl*^ offspring, without a noticeable change in the mRNA expression level of the *Hcn* 1, 3 and 4 isoforms (Figure S1). Consistent with a previous report showing impairment in the growth of mice with constitutive deletion of HCN2 [17], deletion of HCN2 in all neurons resulted in attenuated weight gain (Figure S1). As expected from the sparsity of cholinergic cells throughout the CNS, no changes in the brain expression levels of HCN2 or in the mouse body weight were detected in animals with HCN2 deletion in cholinergic cells (cKO, Figure S1). To assess the effect of the deletion of HCN2 in ChIs on their function, HCN-mediated current (I*h*) was measured in acute brain slices from the medial NAc shell. We found that the amplitude of the HCN current was greatly reduced in ChIs from HCN2 cKO mice (Figure 1A, B, Figure S1). We then examined the role of HCN2 in regulating the cellular response to dopamine (DA) by measuring the changes in the current induced by bath application of DA (1 µM). Application of DA in NAc ChIs from WT mice significantly decreased the HCN current (Figure 1A, B), an inhibitory effect likely to be mediated by the DA D2 receptor, which was highly abundant in these cells (Figure S1). In contrast, DA did not have any effect on the amplitude of the HCN current in ChIs from cKO mice (Figure 1A, B), indicating that HCN2 mediates the cellular response to DA in NAc ChIs.

**Figure 1.**
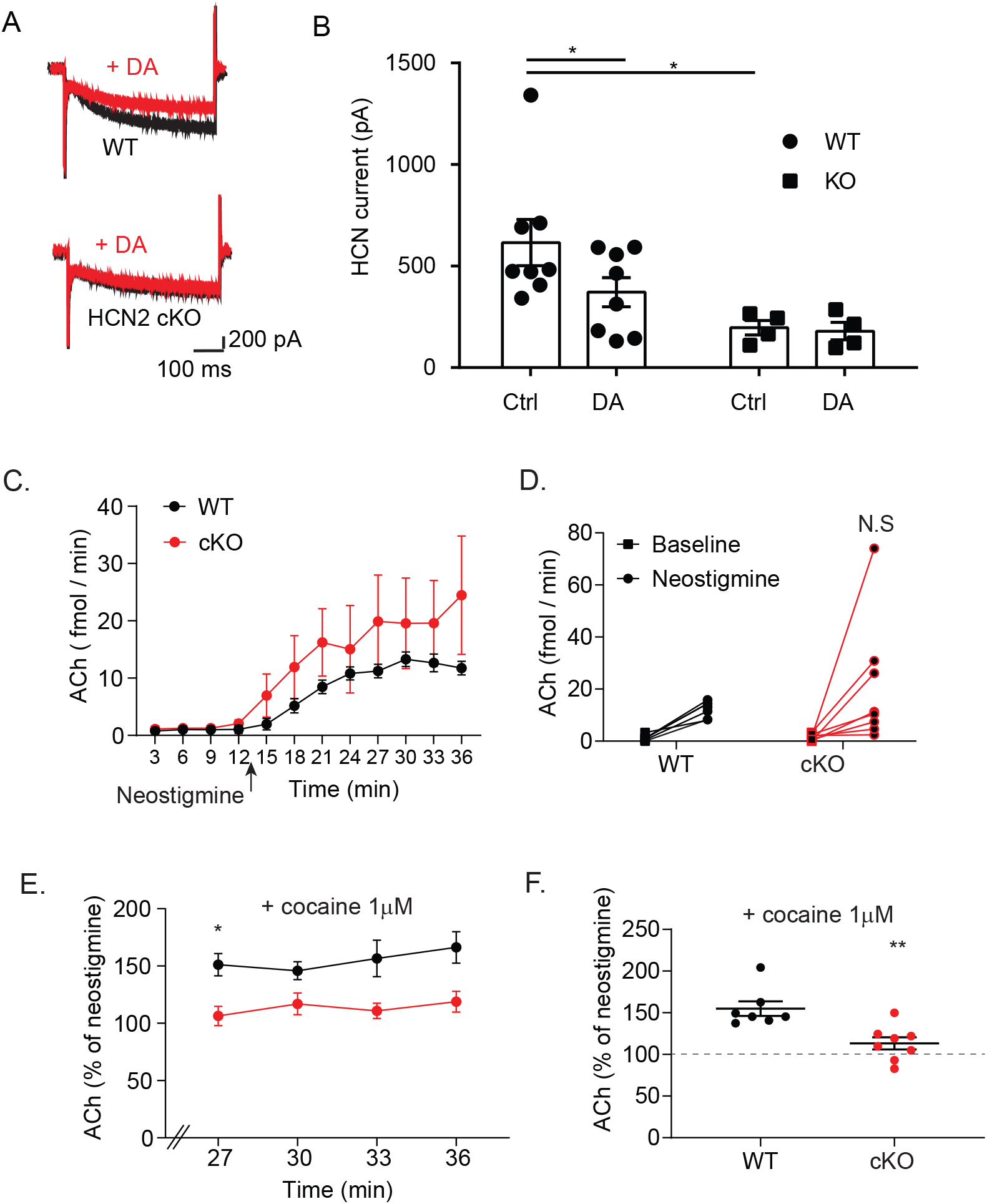
HCN2 regulates cholinergic signaling in the NAc. (A) Representative traces of HCN currents elicited by –150 mV in NAc ChIs from WT and cKO mice before (black, Ctrl) and after DA (1 μM, red). (B) Histograms showing the amplitude of HCN currents at -150 mV in control or DA-treated ChIs from NAc in WT (n = 8 neurons/ 4 mice) and cKO (n = 4 neurons/ 3 mice). Two-way RM ANOVA. Genotype X Treatment *F* [1,10] = 3.39; P= 0.09. Genotype *F* [1,10] = 5.90; P= 0.03. Treatment *F* [1,10] = 4.48; P= 0.06 \**p*< 0.05 by Bonferroni (C) Effect by aCSF without or with neostigmine (500 nM, arrow) on NAc ACh efflux in WT and HCN2 cKO (n= 7, 8 mice, respectively). (D) Means of ACh efflux in the first four bins with aCSF and last four bins with neostigmine. Two-way ANOVA RM. Genotype X Treatment *F* [1,13] = 0.8; P> 0.05. Genotype *F* [1,13] = 1.0; P> 0.05. Treatment *F* [1,13] = 11.33; P= 0.005. N.S. not significant vs. WT. (E) Line graph of ACh efflux during cocaine (1 µM) and neostigmine (500 nM) dialysis in WT and cKO (n= 7, 8), plotted as percentage of that during neostigmine. Two-way RM ANOVA. Genotype X Time *F* [3,39] = 0.65; P> 0.05. Genotype *F* [1,13] = 13.59; P= 0.002. Time *F* [3,33] = 1.29; P> 0.05. \**p*= 0.017 by Bonferroni. (F) Dot plot of means. \*\**p*= 0.003 by unpaired t-test.

We then studied the role of HCN2 in ChIs in regulating acetylcholine (ACh) release, *in-vivo*, by measuring the efflux of ACh in the NAc by microdialysis in freely moving mice. Infusion of the cholinesterase inhibitor, neostigmine (500 nM) elevated ACh efflux in WT and cKO by 1107.3% and 1426.5%, respectively, without a noticeable difference between the two groups (Figure 1C, D). Infusion of cocaine (1 µM) to the NAc further increased ACh efflux by 55.0 ± 8.75% in WT, whereas in cKO this enhancement was initially smaller (113.2 ± 7.88%, Figure 1E, F).

### Dopamine response to reward in the NAc is impaired in HCN2 cKO mice

To study the role of HCN2 in ChIs in regulating DA release from mesolimbic DAergic nerve terminals, the efflux of DA in the NAc was measured in freely moving mice using microdialysis. In line with the fact that the basal efflux of acetylcholine was unchanged in cKO mice, no differences were detected in the basal levels of DA or its metabolites between WT and cKO (Figure 2A). We then measured changes in DA efflux during exposure of freely moving mice to rewarding stimuli. In WT male mice, the efflux of DA was enhanced by exposure to palatable food or to unfamiliar male or female mice, or by local infusion of cocaine (1 or 10 μM, Figure 2B -F). The enhancement in DA release in the NAc in response to exposure to palatable food or female mice was largely attenuated in cKO (Figure 2B, D). Moreover, the enhancement in DA release was delayed in cKO mice following male mouse encounter as well as in mice subjected to local infusion of 1 μM of cocaine (Figure 2C, E), supporting the idea that NAc ChIs are activated by diverse rewarding stimuli. However, high dose of cocaine (10 μM) induced a similar enhancement in the DA efflux between WT and cKO (Figure 2F), suggesting that additional mechanisms contribute to the DA enhancement by high dose of cocaine.

**Figure 2.**
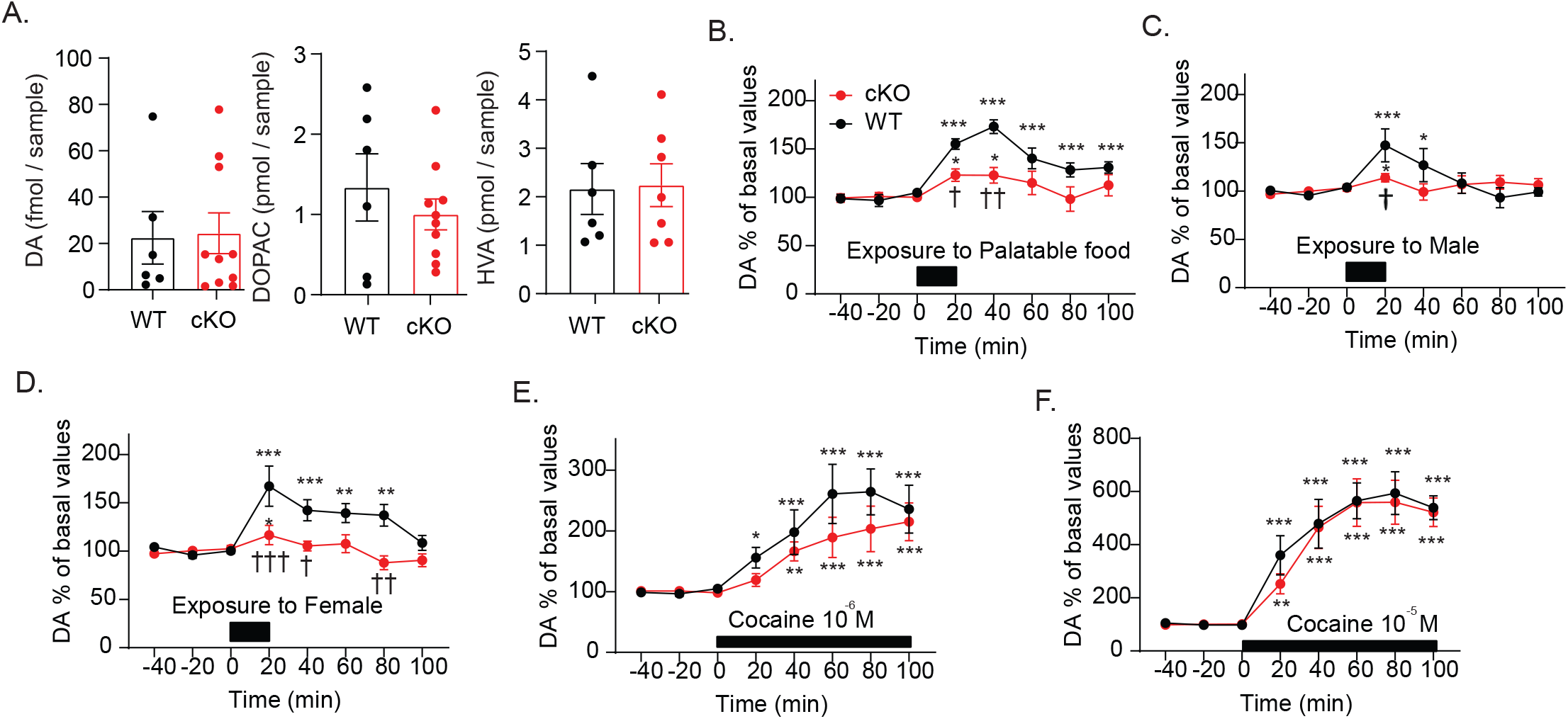
HCN2 in the NAc ChIs mediates the enhancement in DA release by rewarding stimuli. (A) Basal levels of dopamine, DOPAC and HVA in WT and HCN2 cKO mice (n=6, 10 mice, n=6, 7 mice, and n=6, 7 mice, respectively), which were obtained as the average of three stable baseline samples before rewarding stimuli. DA t_(10)_ = 0.1358, *p* > 0.05; DOPAC, t_(7)_ = 0.7281, *p* > 0.05; HVA, t_(10)_ = 0.1164, *p*> 0.05, by Welch’s t-test. (B-D). Responses to exposure to palatable food (B), an unfamiliar male mouse (C), and an unfamiliar female mouse (D) in WT and cKO mice (n = 5, 5 mice, n = 5, 6 mice and n = 5, 5 mice, respectively). The values obtained after rewarding stimuli were compared with the basal levels obtained as the average of three stable baseline samples (100%) using mixed linear models with time as a covariate, and post hoc Bonferroni. \**p* < 0.05, \*\**p* < 0.01, \*\*\**p* < 0.001 vs the basal levels. (B) Two Way ANOVA. Group X Time *F* [7, 64] = 3.04, P= 0.008. Group *F* [1, 64] = 26.57, P< 0.0001. Time *F* [7, 64] = 12.13, P< 0.0001. ^†^*p*= 0.030 ^††^*p*= 0.0001 vs WT by Sidak. (C) Two Way ANOVA. Group X Time *F* [7, 72] = 2.27, P= 0.038. Group *F* [1, 72] = 1.50, P> 0.05. Time *F* [7, 72] = 3.59, P= 0.002. ^†^*p* = 0.033 vs WT by Sidak. (D) Two Way ANOVA. Group X Time *F* [7, 64] = 3.30, P= 0.005. Group *F* [1, 64] = 29.48, P< 0.0001. Time *F* [7, 64] = 6.80, P< 0.0001. ^†^*p* = 0.026, ^††^*p* = 0.001, ^†††^*p* < 0.001 vs. WT by Sidak. (E, F) Responses to NAc infusion of cocaine at the concentrations of 1 μM (E) and 10 μM (F) in WT and cKO mice (n= 5, 5 mice and n=5, 5 mice, respectively). (E) Two Way ANOVA. Group X Time *F* [7, 64] = 0.5866, P> 0.05. Group *F* [1, 64] = 4.54, P= 0.036. Time *F* [7, 64] = 10.82, P< 0.0001. (F) Two Way ANOVA. Group X Time interaction *F* [7, 64] = 0.20, P> 0.05. Group *F* [1, 64] = 0.61, P> 0.05. Time *F* [7, 64] = 29.09, P < 0.0001.

To study how ChIs initiate an enhancement in DA signaling in the NAc by reward, a DA biosensor was applied. AAV5 with the DA biosensor dLight 1.3b was injected into the medial NAc, and optic fiber was implanted to record fluorescent signals in anesthetized mice (Figure 3A). Systemic injection of cocaine (1.5 mg/ kg) induced a 133.5% increase in the photometric response, relative to saline, with peak emission 6.8s post injection (Figure 3B-D). The specificity of this response to dopamine signaling was then validated. Cocaine injection reversed an attenuation in fluorescence by the dopamine D2 receptor antagonist, haloperidol (2 mg/kg, Figure S2), and the photometric response to cocaine was completely inhibited by pre-treatment with the dopamine D1 receptor antagonist, sch23390 (0.2 mg/ kg), or in animals that were not transfected with the dLight 1.3b bio-sensor (not shown).

**Figure 3.**
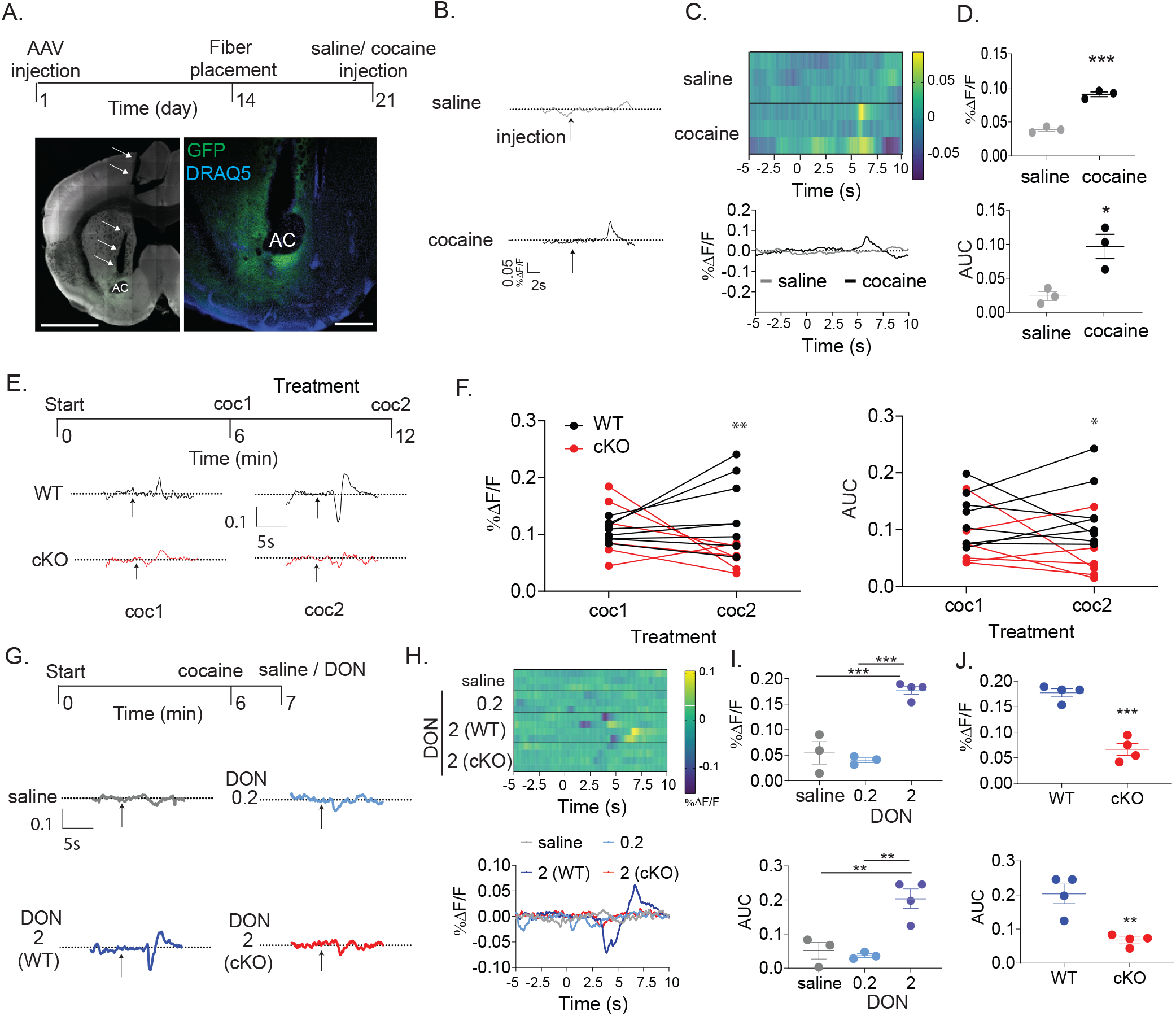
NAc ChIs mediate the enhancement in dopamine signaling by cocaine. (A) Study design and representative microscope images showing the localization of the fiber trace (arrows) and the dLight1.3 GFP in the NAc. AC, anterior commissure. Scale bars, 500 µm. (B) Representative photometric traces between 5s before and 10s after intraperitoneal injection of saline or cocaine (1.5 mg/kg). Arrows indicate injection times, dashed line represents %ΔF/F= 0. (C) Perievent time heatmap of individual samples(top) and histogram of group means (bottom) of saline or cocaine in n= 3 mice / group. (D) Effect of cocaine (1.5 mg/kg) injections in WT (n= 3 mice) on fluorescent peak amplitude (top) and area (AUC, bottom). \**p*= 0.018, \*\*\**p*= 0.0003, vs. saline by unpaired t-test. (E) Study design and representative traces of two consecutive injections of cocaine (1.5 mg/kg) in WT and HCN2 cKO mice. (F) Changes in fluorescent peak amplitude and area by consecutive cocaine injections in WT (n= 8 mice) and cKO (n= 6 mice). Amplitude Two Way ANOVA RM, *F* genotype X treatment [1, 12] = 7.39, P= 0.019. \*\**p*= 0.007 between genotypes by Sidak. Area Two Way ANOVA RM, *F* genotype X treatment [1, 12] = 1.053, P> 0.05. *F* genotype [1, 12] = 6.65, P= 0.024. *F* treatment [1, 12] = 0.53, P> 0.05. \**p*= 0.026 between genotypes by Sidak. (G) Study design and representative traces following injections of cocaine (1.5 mg/kg) followed by saline or donepezil (DON, 0.2 or 2 mg/kg) in WT and HCN2 cKO. (H) Perievent time heatmap and histogram of averages by saline (n= 3, WT), donepezil 0.2 mg/kg (n= 3, WT) or 2 mg/ kg in WT or cKO (n= 4 mice each). (I) Changes in fluorescent peak amplitude and area in WT. One Way ANOVA (amplitude), P= 0.0002, \*\*\**p*< 0.001 by Tukey. One Way ANOVA (area), P= 0.003, \*\**p*< 0.01 by Tukey. (J) Effect of donepezil (2 mg/kg) injections in WT and cKO (n= 4, 4 mice) on fluorescent peak amplitude and area. \*\**p*= 0.004, \*\*\**p*< 0.001 by unpaired t-test.

The response to repeated injections of cocaine (1.5 mg/ kg) was then tested in WT and cKO mice (Figure 3E). A second injection of cocaine induced a response in WT similar in magnitude to that by the first (Figure 3F). In contrast, in cKO mice, the second injection of the drug resulted in a 27.6% reduction in the dopaminergic response relative to that by the first injection, with no difference between the genotypes in their response to the first injection (Figure 3F). To study the possibility that activation of cholinergic signaling by cocaine mediated the enhancement in DA signaling, we administered cholinesterase inhibitor after cocaine (Figure 3G). In WT mice, injection of 2 mg/ kg donepezil, but not 0.2 mg/ kg, induced a dramatic 224.8% increase in the dopaminergic response, relative to saline (Figure 3H, I). In contrast, donepezil had no effect on the DAergic response in cKO mice (Figure 3J), whereas saline injection following cocaine had no effect on either genotypes. Finally, we tested the possibility that the attenuated cholinergic signaling by cocaine in cKO is mediated by changes in nicotinic receptor function. Nicotine (0.07 mg/ kg) induced photometric responses in WT and cKO, with a higher response in the latter, ruling out the possibility that nicotinic receptor function in DAergic neuronal terminals is impaired in cKO mice (Figure S2). Taken together, these data strongly support the idea that NAc ChIs activation by rewards plays a role in mediating the enhancement in DA release from local nerve terminals.

### The behavioral response to reward is attenuated in HCN2 cKO mice

The behavioral implications of this impaired response to reward was then tested in cKO mice. In the open field test, HCN2 cKO mice showed a 64.7% increase in motor activity after a single cocaine injection (15 mg/ kg), relative to WT mice, without any difference between their locomotion in drug-naïve cohort (Figure 4A). In line with the impairment in enhancing DA release in response to social encounter, HCN2 cKO mice showed reduced social approach behavior in stress-naïve environment as well as after mild social stress (Figure S3). To test the motivational effect by cocaine, a conditioned place preference (CPP) paradigm was used [18]. Mice were trained to pair one chamber with cocaine (15 mg/ kg) and one with saline (Figure 4B). In the CPP acquisition test, WT mice, but not cKO, spent 143 ± 31.52% more time in the drug-paired compartment relative to that in the saline-paired chamber. On the contrary, cKO mice spent less time in the drug-paired compartment and more time in the saline paired chamber relative to WT mice (Figure 4C), whereas locomotion during the test was not different between the groups (Figure S3). The behavioral response to cocaine after withdrawal was then tested. No differences were detected between WT and cKO in their acquisition of CPP extinction. In WT mice, cocaine priming after 1 day or 30 days of extinction, induced a 283.8% and 280.9% increase in the respective time spent in the cocaine-associated chamber, relative to that in the saline-associated chamber. In line with the impairment in acquisition of CPP, cocaine priming failed to reinstate CPP in cKO mice (Figure 4C).

**Figure 4.**
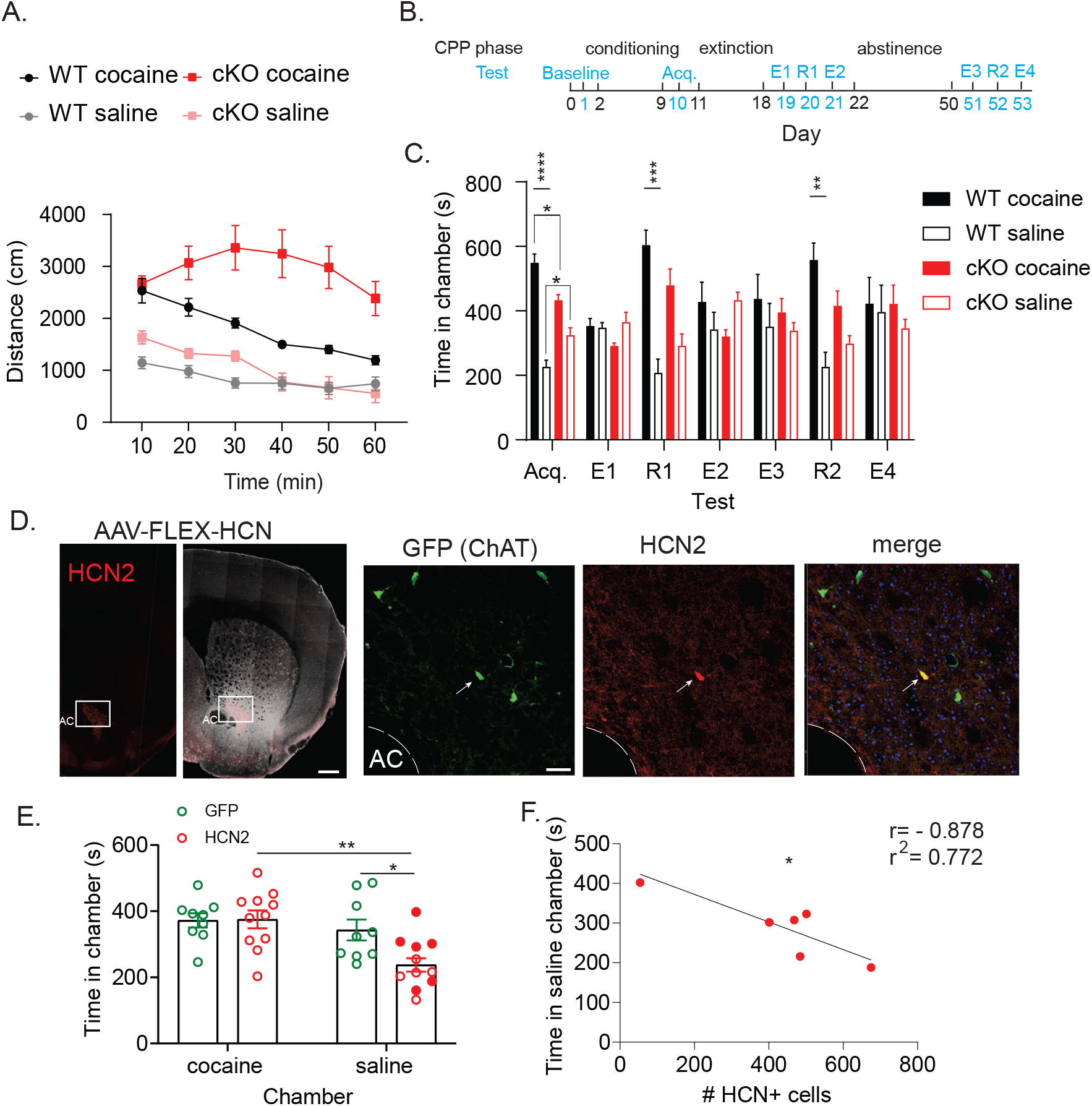
HCN2 in NAc ChIs mediates the acquisition of cocaine CPP. (A) Distance traveled in WT and cKO during 60 min after cocaine (15 mg/kg, n= 6, 5) or saline (n= 8, 7). Cocaine Two Way ANOVA RM, *F* genotype X time [5, 45] = 7.68, P< 0.0001, *F* genotype [1, 9] = 14.57, P= 0.004. *F* time [2, 18] = 9.98, P= 0.001. Saline Two Way ANOVA RM, *F* genotype X time [5, 65] = 4.94, P< 0.0007, *F* genotype [1, 13] = 1.87, P> 0.05. *F* time [3, 37] = 20.92, P< 0.0001. (B) CPP design. Black indicates paradigm phases and their day numbers, and cyan indicates tests names and their day numbers. Acq, acquisition. E, extinction. R, reinstatement. (C) Time spent in each chamber in WT and cKO (n= 8, 5) during the different tests. Two Way ANOVA RM, *F* genotype X phase [18, 128] = 4.96, P< 0.0001, *F* genotype [3, 22] = 4.75, P= 0.010. *F* phase [3, 61] = 0.84, P= 0.476. \*\*\*\**p*< 0.0001,\*\*\**p*< 0.001, \*\**p*< 0.01, \**p*< 0.05 by Tukey. (D) Representative low (left) and high (right) magnification immunohistochemical images following stereotaxic injection of AAV.FLEX.HCN2 in NAc of cKO mouse. Arrows indicate HCN2 labeling in ChI. Scale bars, 500 μm (left) and 50 μm (right). AC, anterior commissure. (E) Time spent in each chamber during CPP acquisition test in cKO mice injected with AAV.FLEX.GFP (n= 9) or AAV.FLEX.HCN2 (n= 11). Two Way ANOVA, *F* AAV X chamber [1, 18] = 5.10, P= 0.037, *F* AAV [1, 18] = 3.73, P> 0.05. *F* chamber [1, 18] = 12.08, P= 0.004. \**p*= 0.011 by Sidak. (F) Correlation between time in the saline-associated chamber and number of HCN2 immunopositive ChIs in cKO mice injected with AAV.FLEX.HCN2 (n= 6, indicated as full red circles in E). \**p*= 0.021 by Pearson.

We then tested whether restoring HCN2 levels selectively in NAc ChIs of cKO mice could correct the deficit in the acquisition of cocaine CPP. The previously validated AAV-FLEX-HCN2 [5] was stereotaxically injected to the NAc of cKO mice, and immunostaining confirmed the expression of HCN2 protein in their NAc ChIs (Figure 3D). Restoring HCN2 in NAc ChIs of cKO resulted in reduced time spent in the saline-associated chamber during the CPP acquisition test, relative to that in the drug-associated chamber (Figure 4E). Immunohistochemical analysis identified an association between the number of HCN2 immunopositive NAc ChIs and the reduction in time spent in the saline-associated chamber (Figure 4F).

We then studied how an increase in cholinergic activity might affect cocaine conditioning. First, we studied the effect of over-expression (O/E) of HCN2 in NAc ChIs on the acquisition, extinction and reinstatement of cocaine CPP. To this end, ChAT ^*Cre*^ mice were stereotaxically injected with either AAV-FLEX-HCN2 or control AAV-FLEX-GFP to the NAc (Figure 5A). Both GFP O/E and HCN2 O/E mice showed preference to the cocaine-associated chamber in the acquisition test, whereas HCN2 O/E mice spent more time in the cocaine-associated chamber and less time in the saline-associated chamber during the reinstatement test, relative to control (Figure 5B).

**Figure 5.**
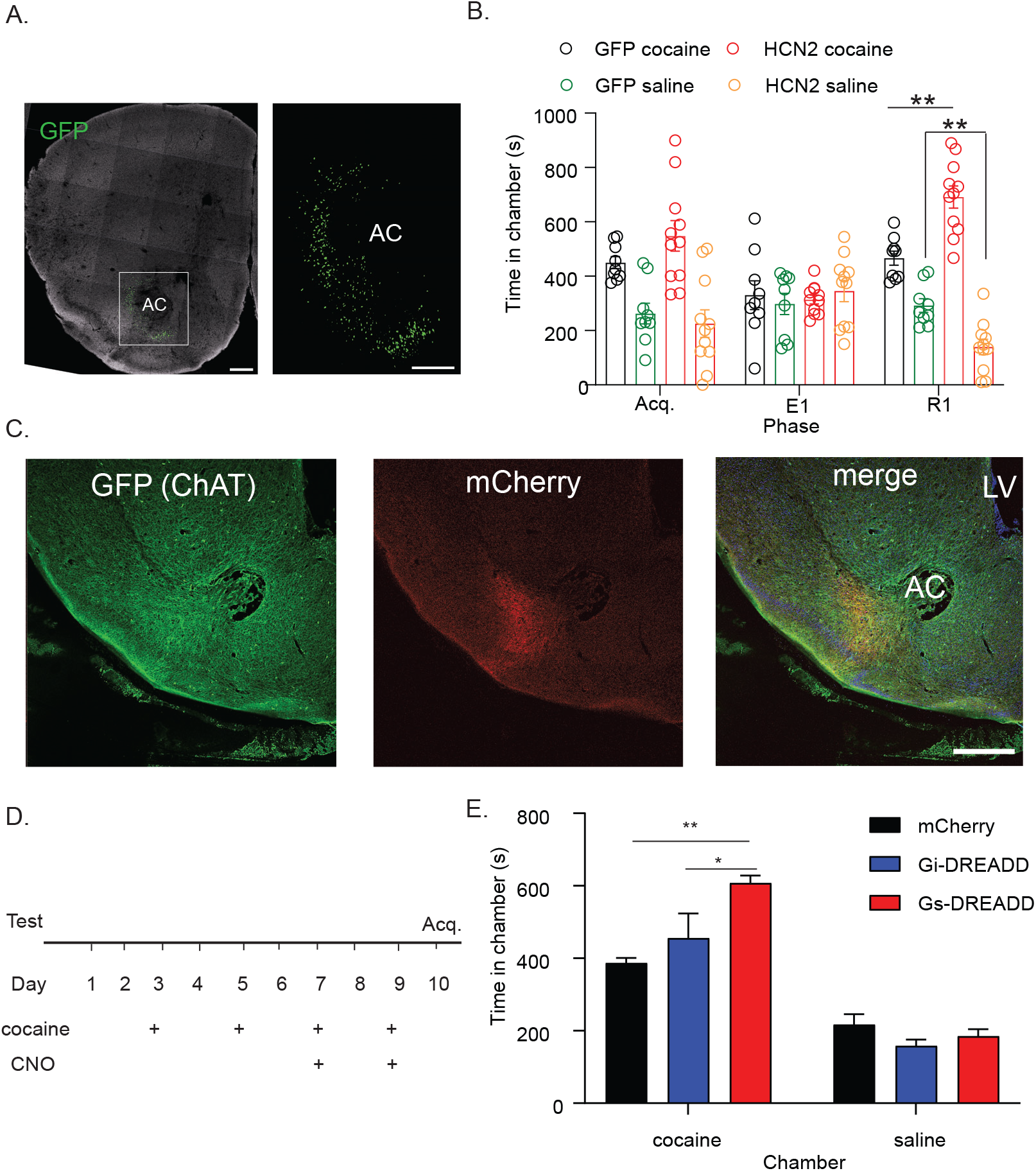
Activation of NAc ChI enhances cocaine CPP. (A) Representative immunohistochemical images depicting GFP labeling following stereotaxic injection of AAV.FLEX.GFP in the NAc of ChAT ^*Cre*^ mouse. Scale bars, 500 μm. AC, anterior commissure. (B) Time spent in the saline- and cocaine-associated chambers during CPP acquisition (Acq) extinction (E1) and cocaine-induced reinstatement (R1) tests in ChAT^*Cre*^ mice injected with AAV.FLEX.GFP (n= 8) or AAV.FLEX.HCN2 (n= 11). Two Way ANOVA, *F* phase X chamber [6, 64] = 5.25, P= 0.0002. *F* phase [2, 71] = 4.58, P= 0.014. *F* chamber [3, 36] = 25.41, P< 0.0001. \*\**p*< 0.01 by Tukey. (C-E) Chemogenetic manipulation of NAc ChIs. (C) Representative immunohistochemical images depicting mCherry labeling following stereotaxic injection of AAV2-hSyn-DIO-mCherry to the NAc of ChAT^*Cre*^ mouse. Scale bars, 500 μm. AC, anterior commissure; LV, lateral ventricle. (D) Study design. CPP conditioning started three weeks after the stereotaxic AAV injection. CNO (4 mg/kg) was injected to all animals on days 7 and 9 of the conditioning phase, 15 min before cocaine. (E) Time spent in the saline- and cocaine-associated chambers during the CPP acquisition test (Acq) in ChAT^*Cre*^ mice injected with Gi-DREADD, Gs-DREADD or mCherry control (n= 5 mice in each group). Two Way ANOVA, *F* genotype X chamber [2, 24] = 7.13, P= 0.0037. *F* genotype [2, 24] = 5.03, P= 0.015. *F* chamber [1, 24] = 118.20, P< 0.0001. \**p*= 0.038, \*\**p*= 0.0012 by Tukey.

To test the effect of activation of NAc ChIs on cocaine CPP, a chemogenetic approach was applied. ChAT ^*Cre +/-*^ mice were bilaterally injected with AAVs expressing inhibitory DREADD (AAV2-hSyn-DIO-hM4D(Gi)-mCherry), excitatory DREADD (AAV2-hSyn-DIO-rM3D(Gs)-mCherry) or a control vector (AAV2-hSyn-DIO-mCherry) to the medial NAc.

Immunohistochemical analysis confirmed that the DREADD viruses infected NAc cells (Figure 5C). To induce a subtle effect on CPP, clozapine-N-oxide (CNO) was injected before two cocaine sessions of the conditioning phase (Figure 5D). During the acquisition test, all groups showed preference to the cocaine-associated chamber, relative to the saline-associated chamber. Notably, activation of ChIs during conditioning increased the time spent in the cocaine-associated chamber by respective 33.3% and 56.9%, relative to that in the groups injected with either Gi-DREADD or mCherry (Figure 5E), suggesting that enhancement of NAc ChIs activity during exposure to rewarding stimuli facilitates reinforcement.

## Discussion

Here we report that the physiological and behavioral responses to diverse rewarding stimuli are mediated by cholinergic neurons of the NAc. Activation of NAc ChIs by rewarding stimuli mediates an enhancement in DA release and in its signaling in the NAc, and is essential for the behavioral response to social encounter and cocaine conditioning. The efflux of acetylcholine in the NAc is elevated by local cocaine infusion, and the physiological and behavioral responses to cocaine are respectively induced by enhancement of cholinergic signaling or by chemogenetic activation of NAc ChIs, highlighting a new role for these cells in reward encoding. The changes in cholinergic signaling by rewarding stimuli depends on HCN2 in ChIs, which mediates the HCN current in ChIs as well as its modulation by DA signaling. HCN2 in ChIs is essential for the behavioral response to reward and represents a novel target for the development of treatment for drug addiction.

### HCN2 regulates the HCN current and DA signaling in NAc ChIs

Deletion of HCN2 in ChIs resulted in attenuation in the HCN current and in its modulation by DA. A role for DA signaling in ChIs function is well established [19]. In the dorsal striatum, DA agonists inhibit ChI activity via D2 receptors activation [20]. DA signaling in ChIs has been shown to down regulate the persistent sodium [21] and HCN currents in a cAMP dependent manner, thus inhibiting tonic firing [22]. In line with this, the presence of a conditioned stimulus associated with reward evokes a pause in the firing of tonically active ChIs [23], [24] In support of the role of dopamine D2 receptors in mediating the effect by DA on HCN current in NAc ChIs, we identified high expression levels of the dopamine D2 receptors in ChIs of both the NAc and the dorsal striatum. Moreover, the physiological and behavioral responses to cocaine were attenuated in HCN2 cKO mice, which is in line with the impaired motivation for self-administration by cocaine following down regulation of D2 receptors in NAc ChIs [25].

Although HCN2 is the most abundant HCN isoform in NAc ChIs, HCN1 and HCN4 are also co-expressed in NAc ChIs [5]. In contrast to the deletion of HCN2 in cKO, the expression of the other HCN isoforms was unchanged, suggesting that the HCN current and its modulation by DA are primarily regulated by the HCN2 homotetramer. An impairment in the HCN current following selective down-regulation of HCN2 in ChIs was suggested after deletion of p11 (s100a10) in ChIs or in stress-sensitive mice [5]. In those mice, the impairment in the HCN current was accompanied by reduced firing rate in tonically active NAc ChIs. Therefore, the impairments we report here in the HCN current and its modulation by DA in ChIs from HCN2 cKO mice, suggest that the HCN2 isoform mediates both the tonic firing in NAc ChIs as well as the pause in their tonic firing by DA signaling activation. In line with this idea, genetic deletion of HCN2 in nociceptive neurons abolished both their enhancement of firing caused by an elevation of cAMP and the inhibitory effect of HCN2 inhibition on firing frequency [26, 27].

### Activation of cholinergic signaling by reward mediates the enhancement in DA release in the NAc

Genetic deletion of HCN2 in ChIs resulted in a specific impairment in both cholinergic and DAergic signaling in the NAc. The expected increase in acetylcholine and DA effluxes following exposure to rewarding stimuli were delayed and attenuated, without a parallel effect on their basal levels. The enhancement in DA efflux by food reward and female encounter was diminished, and that by male encounter or by low dose of locally infused cocaine was delayed. In line with the effect on DA level, HCN2 cKO mice also showed impairment in the enhancement in DA signaling by systemic administration of cocaine or cholinesterase inhibitor. The enhancement in DA efflux in the NAc by rewarding stimuli is required for the behavioral response to reward [28], but a mechanism by which cholinergic signaling is enhanced by reward has not received much attention. We previously found that chemogenetic inhibition of NAc ChIs attenuated DA release in response to cocaine, whereas chemogenetic activation enhanced it [8]. This is in line with the facts that the spontaneous release of DA from NAc nerve terminals is mediated by ChI activity [2], and that the reward response in tonically active ChIs consists of an initial excitation, a pause and rebound excitation [29]. Other firing patterns of ChIs including irregular discharge and rhythmic burst firing [30], could also contribute to the cholinergic enhancement by reward, as ChIs also show transition to bursting activity by activation of DAergic signaling [31]. Such bursts increase the reliability of information transfer, especially in the presence of unreliable synapses [32], such as those by the dense axonal arborization of ChIs onto DAergic nerve terminals. In addition to the potential increase in autonomous activity, changes in inhibitory and excitatory signaling in ChIs could also contribute to the activation of NAc ChIs by reward. Reward response activates glutamate co-release onto ChIs [33], whereas inhibitory signaling from VTA inhibitory neurons is likely attenuated [34, 35].

Two distinct mechanisms could mediate the enhancement in DA release in the NAc by rewarding stimuli. DAergic cell firing or activation of receptors at their axonal terminals can each enhance DA transmission in the NAc. In support of the idea that the activation of ChIs is likely to contribute to local enhancement in DA signaling in response to reward, the release of DA in the NAc by rewarding stimuli does not correlate with the activity of VTA DA cells, suggesting that motivation-related DA release in the NAc does not arise from DA cell spiking [36]. Among NAc cell types, ChI activity directly enhances DA release from terminals [3], whereas that of spiny projection neurons indirectly regulate DAergic cell activity. In support of the idea that ChI activity enhances local DA release during exposure to rewarding stimuli, without affecting DAergic cell activity, an impairment in NAc ChI function did not affect the physiological response to reward in other circuits. Deletion of p11 in ChIs attenuated the increase in DA in response to cocaine in the mesolimbic (VTA-NAc) but not the mesocortical (VTA-PFC) projection pathway [8]. Our present work suggests that the interaction between the activities of DAergic cells and ChIs is dynamic and changes after prolonged exposure to rewarding stimulus or by its magnitude. The enhancement in DA levels by 1 μM cocaine was detected after 20 min of exposure in WT whereas that in HCN2 cKO mice was detected only after 40 min, and the differences between WT and cKO in their responses to other rewarding stimuli were most notable immediately after the exposure to the stimulus. It is therefore likely that the initial exposure to rewarding stimuli activates ChIs, whereas prolonged exposure inhibits them. This is in line with the fact that an increase in DAergic cell firing by cocaine is delayed [37]. Moreover, the enhancement in DA level by exposure to higher dose of cocaine, which mimics high rate of DAergic cell activity was not attenuated in cKO mice, suggesting that NAc ChIs activation contributes to reward encoding in the absence of extensive DAergic cell firing. In light of our findings, it is suggested that the enhancement in cholinergic signaling by rewarding stimuli is required to initiate the reward response, whereas the delayed activation of DAergic neurons and subsequent inhibition of ChIs activity mediates a delayed phase of this response.

### HCN2 in NAc ChIs regulates reward encoding and pleasure response

In line with the idea that HCN2 in ChIs initiate the reward response by activating these cells, chemogenetic activation of NAc ChIs during conditioning enhanced the time spent in the cocaine-paired chamber during the acquisition test. The idea that activation of ChIs mediates the behavioral response to reward is in line with previous reports showing that optogenetic inhibition of NAc ChIs impaired cocaine conditioning, whereas their chemogenetic inhibition impaired the response to food reward and social encounter [4, 5]. Interestingly, an inhibitory response in ChIs to DA and serotonin signaling seem to also play a role in mediating the behavioral response to reward [38],[13, 25], further supporting the idea that dynamic changes in ChI activity are equally important for reward encoding.

The deletion of HCN2 in cholinergic cells resulted in a notable behavioral impairment in the response to reward. The acquisition and reinstatement of cocaine-induced CPP were impaired without a change in the acquisition of extinction. The deficit in conditioning was reversed by restoring HCN2 levels in NAc ChIs. Moreover, up-regulation of HCN2 in NAc ChIs enhanced the time spent in the cocaine-paired chamber and reduced that in the saline-paired chamber during cocaine-induced reinstatement test, indicating that the levels of HCN2 in NAc ChIs dynamically controls conditioning in response to reward. In line with a role for HCN2 in regulating the DA signling, attenuation of DA D2 receptors in NAc ChIs impaired the susceptibility to cocaine seeking [25]. We previously found that the expression level of HCN2 in ChIs is dynamically regulated by the emotional state. HCN2 mRNA level was down regulated in NAc ChIs in models of depression, including stress-sensitive mice following chronic social defeat paradigm as well as in p11 cKO [5]. Stress-sensitive mice and p11 cKO mice showed anhedonia-like behaviors that included reduced sociability, helplessness and impairment in approaching food rewards [5, 6, 13]. Together, our previous and present work suggest that an elevated level and activity of HCN2 in the NAc shell ChIs enable reward encoding whereas its absence leads to impairment in reward response and anhedonia. In light of these findings, it is proposed that HCN2 in NAc ChIs could be a viable target for the development of a cell-type specific treatments for mood disorders including depression and drug abuse.

## Supporting information

Supplemental information, Lee et al.

## Acknowledgements

We are grateful to Adam Ogilvie for proofreading the manuscript. This work was supported by the United States Army Medical Research Acquisition Activity Grant W81XWH-14-0130 (Y.S.), the Fisher Center for Alzheimer’s Research Foundation (M.F), grant-in-Aid for Scientific Research from the Japan Society for the Promotion of Science 19H03410 (A.N.) and the NCATS/NIH-CTSA UL1 TR001866 (Y.S.).

## Author contribution

Conceptualization, YS; Investigation: JL, MW, YK, GU, LM, and YS. Writing – Original Draft, YK, LM, MF, AN and YS, Resources, YS.; Supervision, AN and YS.

## Notes

### Competing Interest Statement

The authors have declared no competing interest.

