## Supplemental information, Lee et al. for "HCN2 in cholinergic interneurons of the nucleus accumbens mediates reward response"

**Legends to Supplemental Figures**

**Fig S1. Validation of the HCN2 cKO mice (related to Fig. 1).**

(A) Bar graph summary of qPCR analysis of *Hcn1-4* mRNA expression in the NAc of: ChAT $^{Cre+/-}$ :: HCN $^{fl/fl}$ (ChAT, cKO), Nestin $^{Cre+/-}$ :: HCN $^{fl/fl}$ (Nes), Nestin $^{Cre-/-}$ :: HCN $^{fl/fl}$ (fl, n= 3 mice per group). ****$p< 0.0001$ by Tukey. (B) Western blot scan showing that HCN2 protein is undetected in NAc protein samples from Nestin $^{Cre+/-}$ :: HCN $^{fl/fl}$ (Nestin) mice. Numbers represent protein size in kDa. (C) Reduced body weight in 8-week old Nestin $^{Cre+/-}$ :: HCN $^{fl/fl}$ (n= 5 male and 5 female), relative to ChAT $^{Cre+/-}$ :: HCN $^{fl/fl}$ (n= 4 male and 5 female). ***$p< 0.001$ by unpaired t-test. (D) Representative traces of HCN currents in ChIs from NAc in WT and mice with HCN2 deleted in ChAT cells (HCN2 cKO), obtained by holding the membrane potential from -60 to -150 mv in 10 mV steps. (E) Bar graph summary showing the amplitude of the HCN current at -150 mV in ChIs from NAc shell in control and HCN cKO mice (n= 4 neurons / 3 mice for each genotype). *$p< 0.05$ by unpaired t-test. (F) Expression level of DA receptors in ChIs. Translated mRNA was isolated using translating ribosome affinity purification from ChIs of the NAc (n= 5 samples / 5 mice) and dorsal striatum (dSt). *$p = 0.032$ vs NAc by unpaired t-test.

**Fig S2. Effect of haloperidol and cocaine on DA biosensor activity (related to Fig. 3).**

(A) Representative photometric traces of 10 min recordings of untreated animal (Baseline, grey), or animal injected with haloperidol (Halo, 2 mg/kg) followed by cocaine (coc, 15 mg/kg). Note that the initial increase in fluorescent signal by haloperidol is likely mediated by inhibition of D2 autoreceptors. (B) Representative traces following injection of nicotine hydrogen tartarate (0.07 mg/kg) in WT and HCN2 cKO. (C) Perievent time heatmap and histogram of averages by nicotine. (D) Effect of nicotine hydrogen tartarate (0.07 mg/kg) in WT and cKO (n= 6, 5 mice) on fluorescent peak amplitude and area. ***p*= 0.008 by unpaired t-test.

**Fig S3. Effect of HCN2 in ChIs on sociability (related to Fig. 4).**

(A) Duration and number of visits in the social approach test in WT (n= 9) and cKO (n= 10). **p*= 0.028, ***p*= 0.003 vs. WT by unpaired t-test. (B) Time ratio (left) and total time spent (right) in the interaction zone (IZ) in WT (n= 18) and cKO (n= 20). **p*< 0.05 vs. WT by unpaired t-test. (C) Beam breaks in each chamber in WT and cKO (n= 8, 5) during the CPP acquisition test. Two Way ANOVA *F* genotype X chamber [1, 22] = 0.58, P> 0.05, *F* genotype [1, 22] = 0.69, P> 0.05. *F* chamber [1, 22] = 16.11, P= 0.0006.

Figure S1

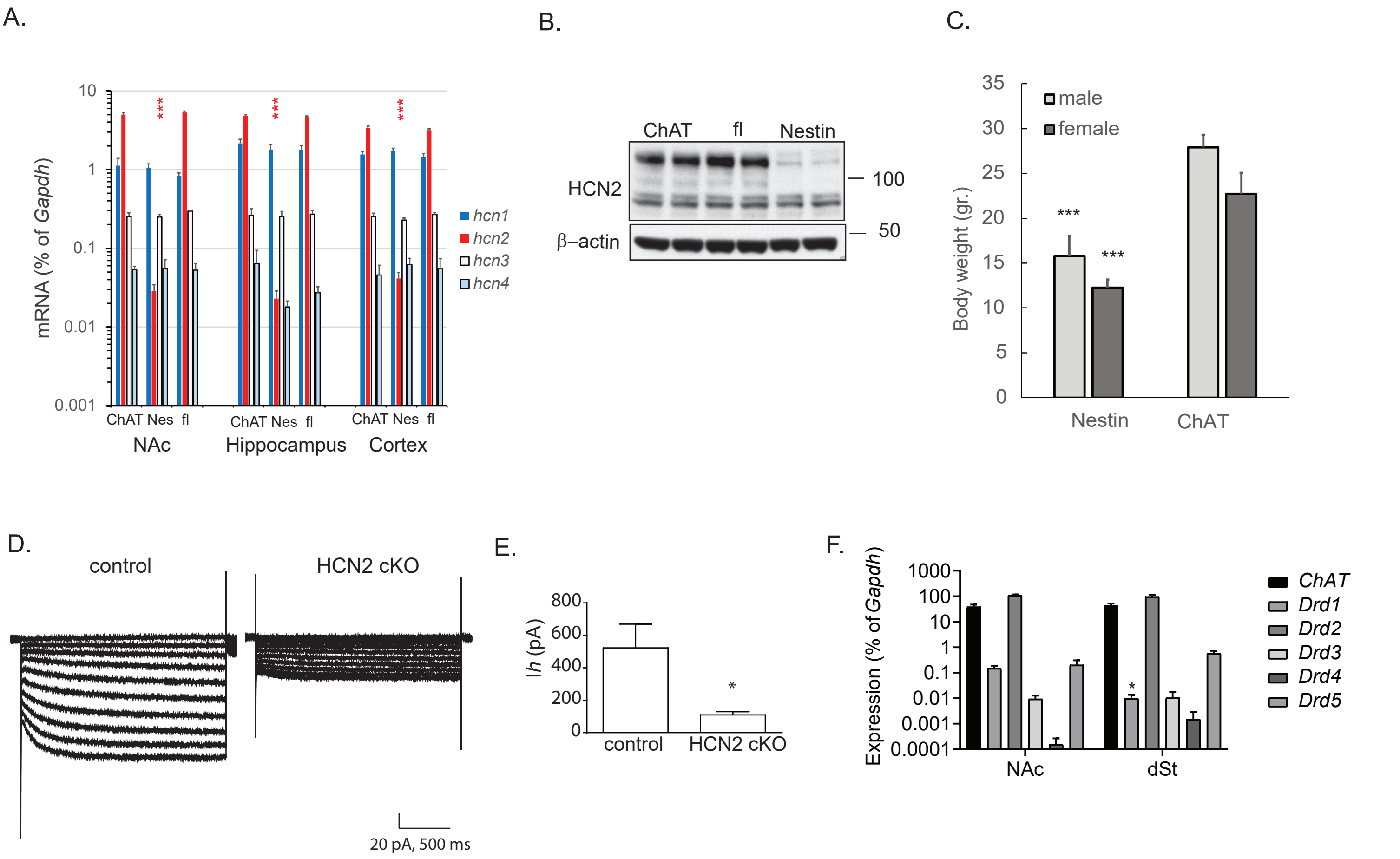

A.
mRNA (% of Gapdh)
10
1
0.1
0.01
0.001
***
***
***
hcn1
hcn2
hcn3
hcn4
ChAT Nes fl
ChAT Nes fl
ChAT Nes fl
NAc
Hippocampus
Cortex
B.
ChAT
fl
Nestin
HCN2
β–actin
100
50
C.
35
30
25
20
15
10
5
0
male
female
Body weight (gr.)
***
***
Nestin
ChAT
D.
control
HCN2 cKO
20 pA, 500 ms
E.
800
600
400
200
0
Ih (pA)
*
control
HCN2 cKO
F.
Expression (% of Gapdh)
1000
100
10
1
0.1
0.01
0.001
0.0001
*
NAc
dSt
ChAT
Drd1
Drd2
Drd3
Drd4
Drd5

### Figure S2

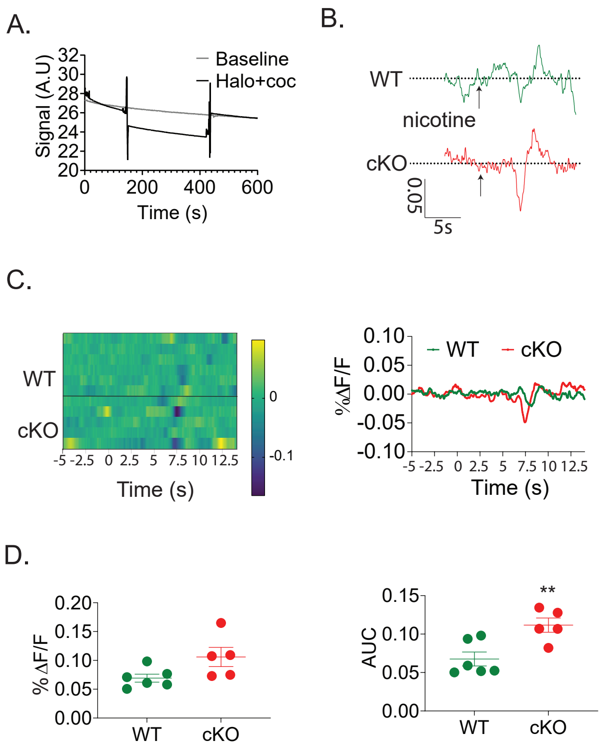
A.
B.
Signal (A.U)
32
30
28
26
24
22
20
0
200
400
600
Time (s)
Baseline
Halo+coc
WT
nicotine
cKO
0.05
5s
C.
WT
cKO
-5 -2.5 0 2.5 5 7.5 10 12.5
Time (s)
0
-0.1
0.10
0.05
0.00
-0.05
-0.10
%ΔF/F
WT
cKO
-5 -2.5 0 2.5 5 7.5 10 12.5
Time (s)
D.
% ΔF/F
0.20
0.15
0.10
0.05
0.00
WT
cKO
AUC
0.15
0.10
0.05
0.00
**
WT
cKO

Fig. S3

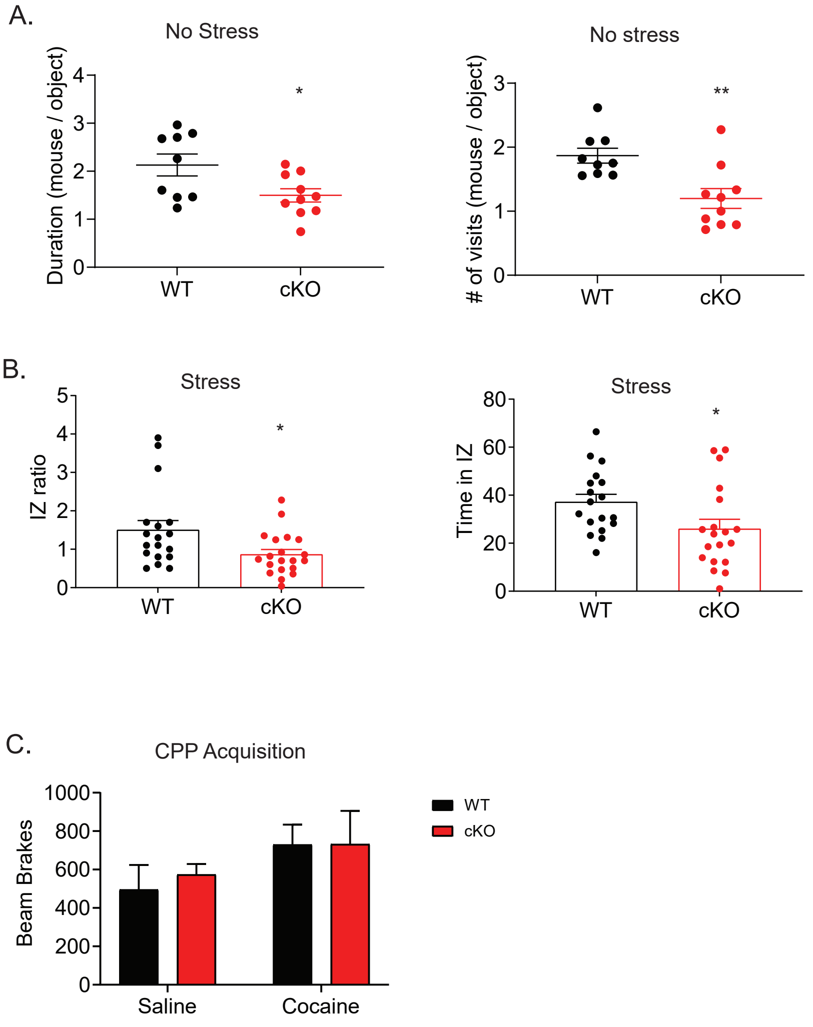
A.
No Stress
Duration (mouse / object)
0
1
2
3
4
WT
cKO
*
No stress
### of visits (mouse / object)
0
1
2
3
WT
cKO
**
B.
Stress
IZ ratio
0
1
2
3
4
5
WT
cKO
*
Stress
Time in IZ
0
20
40
60
80
WT
cKO
*
C.
CPP Acquisition
Beam Brakes
0
200
400
600
800
1000
Saline
Cocaine
WT
cKO
